# Localisation of corticosteroids in male mouse kidney by mass spectrometry imaging

**DOI:** 10.64898/2026.01.16.699839

**Authors:** Ioannis Stasinopoulos, Shazia Khan, Shaden Melhem, Martin Rouamin, Diego F Cobice, Natalie ZM Homer, C Logan Mackay, Roger W Brown, Matthew A Bailey, Ruth Andrew

## Abstract

Renal sodium balance is important for blood pressure homeostasis and is regulated by corticosteroids, chiefly aldosterone but also glucocorticoids. Abundance of these hormones in functional subregions of the kidney is unknown, previous work being limited to measurements in plasma and urine, and in microdialysate from kidney medulla. Here mass spectrometry imaging (MSI) was applied to map corticosteroids in kidney to understand how their distribution overlays with functional targets. Male C57BL6 mice, aged 10 weeks, were fed diets containing 0.03%, 0.3% or 3% w:w sodium for two weeks and kidneys harvested at cull. Steroids were localised as Girard T derivatives on cryosections using Matrix assisted laser desorption ionisation Fourier transform ion cyclotron MSI, with confirmation by liquid extraction surface analysis. Co-registration was performed on sections stained with hematoxylin and eosin. Corticosterone localised along the papilla, medulla and inner cortex, whereas 11-dehydrocorticosterone was concentrated in the medulla. Higher amounts of aldosterone were present in medulla and outer cortex. Distribution patterns were unchanged by dietary salt, but amounts of corticosterone were elevated, particularly in outer cortex with low salt diet. MSI holds great promise to dissect corticosteroid signalling in functional zones of the kidney.

## Introduction

Renal sodium excretion is a key factor regulating sodium balance, which in turn influences extracellular fluid volume and blood pressure. Sodium balance is tightly regulated by the renin-angiotensin-aldosterone axis. Conventionally this is understood to occur via aldosterone activation of mineralocorticoid receptors (MR; NR3C2) in the distal nephron to activate the epithelial sodium channel (ENaC) and stimulate sodium reabsorption. However, cloning of MR and its expression in cells showed that MR was not intrinsically selective for aldosterone and had equal, or perhaps greater affinity for glucocorticoids. (1) Pivotal studies in the 1980s and 90s showed that the enzyme 11β-hydroxysteroid dehydrogenase type 2 (11βHSD2) controls ligand access to MR. This enzyme catalyses rapid conversion of the active glucocorticoid cortisol in humans to inert cortisone (corticosterone to 11-dehydrocorticosterone in mice). (2) The physiological importance of this biochemical control mechanism is shown by the Syndrome of Apparent Mineralocorticoid Excess caused by null mutations in *HSD11B2*, presenting with hypertension that can be lowered by hypothalamic-pituitary-adrenal suppression. (3) Mouse models with reduced or complete loss of enzyme activity show salt-sensitive hypertension due to over activation of ENaC in the distal nephron. (4, 5)

The situation is however more complex with recognition that glucocorticoids can regulate renal sodium transport by activating glucocorticoid receptors (GR; NCR3C1) in the proximal tubule and loop of Henle (6) as well as MR in early stretches of the distal tubules which are unprotected by 11βHSD2. (7, 8) Glucocorticoid excess also causes salt-sensitive hypertension and studies in mice link this to reduced renal salt excretion (9) mediated through MR- and GR-stimulation causing ENaC overactivity. Indeed, GR-mediated control of the thiazide-sensitive NaCl transporter (9) is physiologically important for the circadian rhythm of blood pressure. (10). Type 1 11βHSD adds a further layer of regulation. It is a reductase and is distributed differently throughout nephrons from the type 2 isozyme. It reactivates glucocorticoids where it is expressed e.g. proximal and distal convoluted tubules, renal vasculature, podocytes, macula densa, and interstitial cells of the medulla (6). This complex situation makes it hard to predict whether GR or MR activity will dominate at specific sites and indeed which steroid is the main driver.

Previous measurements of plasma or urinary corticosteroid levels have not allowed regional hormonal activities in tissues to be assessed. Usa et al (11) moved the field forward by interstitial microdialysis of the renal cortex and medulla in mouse suggesting more inactive 11-dehydrocorticosterone than corticosterone in both regions. However, this sampling approach reflects extracellular fluid as opposed to tissue segments. Edwards et al (12) used autoradiography to study binding of tritiated corticosterone and aldosterone in the absence and presence of an inhibitor of 11βHSD2, yielding specific spatial patterns of bound active steroid. However, these data reflect binding to receptors or enzymes and not the available pools of steroids.

Previous studies by our group have utilised Mass spectrometry imaging (MSI) complexed to Matrix assisted laser desorption ionisation (MALDI) to map steroids in brain (13, 14) and testes. (15) Here we adapt MSI to investigate the distribution of corticosteroids in mouse kidney.

## Materials and Methods

### Materials

Girard T (GirT), α-cyano-4-hydroxycinnamic acid (CHCA), trifluoroacetic acid (TFA) and ammonium fluoride (99.99%) were from Sigma-Aldrich (MO, USA). HPLC and LC-MS grade solvents were from Fisher Scientific (Loughborough, UK). Steroids were from Steraloids Inc, PA, USA) and internal standards, D_8_-corticosterone and D_8_-aldosterone (both 2,2,4,6,6,17,21,21-[^2^H_8_]), from Cambridge Isotopes (MA, USA). Superfrost slides were from Thermo Fisher Scientific (MA, USA) and indium tin oxide (ITO) coated slides from Bruker Daltonics (MA, US).

### Animal Protocols

Adult male C57BL/6J mice (Harlan Olac Ltd., Bicester, UK), aged 10-12 weeks, were housed under controlled conditions of temperature (24±10 °C), humidity (50±10%) and light (on 7am to 7pm) with *ad libitum* access to water and commercial rodent chow. Sodium and potassium content of diets (SDS Diets, UK) by weight were respectively: control (0.3%, 0.7% RM1), low salt (0.03%, 0.7%) and high salt (3%, 0.6% w/w). Mice were randomised to treatment groups for 2 weeks. Animals were euthanised (9 to 11 am) by decapitation and trunk blood harvested in sodium citrate 3.2% coated tubes and plasma prepared by centrifugation (5 mins, 14,000 g, room temperature (18-22 °C)). Kidneys were immediately flash frozen in liquid nitrogen and stored at −80 °C. All procedures conformed to the 2010/63/EU Directive and the UK Animals (Scientific procedures) Act, with local ethical review. Adrenal glands from a male Sprague Dawley rat (4-6 months) were harvested into liquid nitrogen as above, following cervical dislocation.

### Sample Preparation

Kidneys and adrenals were supported on 10% w/v gelatin and cryosections (10 µm, −20 °C; HM 525NX cryostat (ThermoScientific, Hemel Hempstead, UK)) mounted on ITO slides. Adjacent sections were harvested onto Superfrost slides for histological analysis. Slides were stored at −80 °C and placed under vacuum (∼20 min) to dry immediately prior to use. Sample preparation was performed with the following modifications. Derivatising reagent and matrix were applied with an automatic sprayer (TM M3 Sprayer (HTX Technologies, NC, USA)) which was cleaned before use by infusing 50% v/v methanol in load and spray modes. GirT derivatisation solution (6 mL; 5 mg/mL in methanol; 0.2% v/v TFA; 6 ng D_8_-corticosterone) was sprayed on the slide (∼50 min): temperature (30 °C), flow rate (0.01 mL/min), criss-cross spraying (15 passes), track spacing (1 mm), velocity (1200 mm/min), pressure (10 psi), gas flow (2 L/min), drying time (2 s), nozzle height: (40 mm). After spraying, slides were incubated in an oven (75% humidity, 40 °C, 1 h) and then placed in a vacuum chamber (20 min). The sprayer was cleaned as above. CHCA matrix solution [10 mL; 10 mg/mL; 50% v/v acetonitrile; 0.2%v/v TFA] was sonicated and sprayed: temperature (85 °C), flow rate (0.24 mL/min), criss-cross spraying (2 passes), track spacing (1 mm), velocity (1200 mm/min), pressure (10 psi), gas flow (2 L/min), drying time (2 s), nozzle height (40 mm). Slides were wrapped in aluminum foil and placed under vacuum until analysis. For LESA-FTICR, tissues were prepared on Superfrost slides and GirT derivatisation performed without matrix application.

For comparison of signal intensities, corticosterone, 11-dehydrocorticosterone and aldosterone standards (1 µg/µL each in methanol) were placed in a glass vial and dried under a stream of nitrogen. GirT derivatisation solution (50 µL; 5 mg/mL in methanol; 0.2% v/v TFA) was added before incubation in an oven (40 °C, 1 h). After derivatisation, solvent was dried under a stream of nitrogen. Derivatised standards were resuspended in CHCA matrix solution (20 µL; 10 mg/mL; 50% v/v acetonitrile; 0.2% v/v TFA) and spotted onto a MALDI steel plate for analysis.

### Sample Analysis

MALDI Fourier transform ion cyclo-tron resonance MS ((FT-ICR-MS) imaging was performed using a 12T SolariX instrument (Bruker Daltonics, Bremen, Germany), with a Smartbeam (1 kHz) laser and operated with SolariX control v2.2.0 (Build 150) and Flex-Imaging v5.0 (Build 80). Instrument parameters were: plate offset (100 V), deflector offset (180 V), laser power (7%, adjusted to achieve maximum sensitivity), laser shots (700, random Smart Walk Pattern), laser frequency 1 kHz, 75 µm raster. For comparison of signal intensities on the MALDI steel plate, the instrument parameters were as follows: plate offset 100 V, deflector offset 200 V, laser power 40%, laser shots 1000 (random smart walk pattern), laser frequency 2 kHz. For targeted analysis, ion scanning was conducted in continuous accumulation of selected ions (CASI) mode with a defined *m/z* 458.30, isolation window *m/z* 50.00 and accumulation time 0.2 s and. For LESA, methanol (1 μL, 50%v/v) was dispensed and retrieved by the LESA source (Advion, Ithaca, NY, USA) and acquired in MS/MS mode using as above, collision induced dissociation (CID); 27 V corticosterone and 11-dehydrocorticosterone, 19 V aldosterone derivatives. Gir T derivatives of corticosterone, 11-dehydrocorticosterone and aldosterone were detected (*m/z* 460.3166 (C_26_H_42_N_3_O_4_), 458.3010 (C_26_H_40_N_3_O_4_), and 474.2957 (C_26_H_40_N_3_O_5_) respectively) and spectral normalisation achieved using D_8_-corticosterone, *m/z* 468.3668. For untargeted markers, data were collated by MALDI-FT-ICR as above in positive mode (Broadband) with range *m/z* 200-2000 without CASI mode. Here, a matrix ion *m/z* 439.029) was used for spectral calibration.

#### Histological Analysis

Tissue (adjacent sections) were rehydrated through a decreasing series of alcohol solutions (100, 95, 80, 70% v/v; 20 sec each) and then the slide was submerged in haematoxylin for 5 mins. It was washed with tap water, submerged in acid-alcohol solution (5-10 sec), washed again and then submerged in ‘Scotts tap water’ solution (20-30 sec). The slide was washed again with tap water, submerged in eosin (5-10 sec), washed and dried and then submerged in increasing series of alcohol solutions (70, 80, 95, 100% v/v; 20 sec each). Finally the slide was submerged in 100% xylene solution (two x 5 mins) and then transferred into pre-mounting xylene solution until mounting. Sections were scanned x40 (Zeiss AxioScan, Jena, Germany). Slides previously subject to MSI were washed twice in 70% v/v ethanol (1 min) to remove matrix and dried prior to staining as above.

### Liquid Chromatography Tandem Mass Spectrometry (LC-MS/MS)

Plasma (50 μL) was defrosted, vortexed thoroughly and centrifuged (2000 g, 10 min, 20 °C; Eppendorf well collection plates, Hamburg,) and diluted with water (50 μL). Internal standards (20 μL) of D_8-_corticosterone (for corticosterone and 11-dehydrocorticosterone) and D_8_-aldosterone (each 5 μg/mL in methanol) were added to each well.

Plasma samples and a 11-point standard curve (0.01-25 ng/mL for 11-dehydrocorticosterone and aldosterone and 10x higher for corticosterone were prepared, Plates were sealed (VWR film, Pennsylvania, USA) and shaken (Microplate Mixer, 5 mins, 600 rpm; Starlab, Milton Keynes, UK). Seals were removed and plates placed in an Extrahera liquid handling robot (Biotage, Uppsala, Sweden), which loaded formic acid (100 μL, 0.1% v/v) into each well. After 5 mins, plate contents were transferred to SLE+ 200 plates (Biotage) before applying positive pressure. Dichloromethane: isopropanol (300 μL, 98:2) was loaded twice into each well and eluate collected in 2 mL deep well plates (Waters, Wilmslow, UK) under positive pressure (0.9 mL extract). Collection plates were reduced to dryness (SPE Dry Argonaut 96 Dual Sample Concentrator, Biotage; gas temperature 40 °C) and residues dissolved in methanol (100 μL, 30% v/v). Plates were sealed (Zone-free sealing film, Waters), shaken (10 min, 600 rpm) and subject to centrifugation (10 min, 4000 g, room temperature).

Extracts containing steroids were analysed by LC-MS/MS according to Lahti-Pulkinnen (16). Steroids were separated on a Kinetex C18 150 × 2.1 mm. 2.6 μm column at 50 °C (Phenomenex, Macclesfield, UK), with an injection volume of 20 μL, using an Acquity I-Class UPLC (Waters). Flow rates were 0.3 mL/min and mobile phase A: H_2_O +0.05 mM ammonium fluoride and B: methanol + 0.05 mM ammonium fluoride followed the gradient: 0-4 min 50-50 A-B; 4-9 min 25-75 A-B; 9-10 min 25-75; 10-12 min 0-100 A B; 12-12.1 min 0-100 A-B; 12.1-16 min 50-50 A-B. Detection was carried out on a QTrap 6500+ (Sciex, Warrington; RRID:SCR_021831) operating in polarity switching electrospray ionisation mode (600 °C, source gas 1 40, source gas 2 60, entrance potential ±10 V, ion spray voltage ±4.5 kV) with optimised declustering potential (DP), collision energy (CE) and collision exit potential (CXP). MRM Transitions monitored were: corticosterone *m/z* 347.1-121.1 (DP=76, CE=29, CXP=8 V), 11-dehydrocorticosterone *m/z* 345.1-121.2 (DP=66, CE=31, CXP=12 V), D_8-_corticosterone *m/z* 355.3-125.1 (DP=56, CE=31, CXP=8 V) in positive mode, and aldosterone *m/z* 359.1-188.9 (DP=−70, CE=−24, CXP=−21 V) and D_8-_aldosterone *m/z* 367.1-193.9 (DP=−75, CE=−26, CXP=−21V) in negative mode. Peak areas were integrated (MultiQuant v3.0.3; Sciex, Warrington, UK; MQ4 algorithm), and amounts interpolated from calibration curves of peak area ratios of steroid:internal standard vs amount, using 1/x weighted regression lines. The method calibration ranges were; aldosterone 0.05–2.5 ng/mL, corticosterone 0.05–250 ng/mL and 11-dehydrocorticosterone 0.10–25 ng/mL. Lower limits of Quantitation (LLOQ) for each steroid in gold serum were 0.05 ng/mL for aldosterone (4.5% intra-assay, 5.5% inter-assay %CV), 0.05 ng/mL for corticosterone (7.9% intraassay, 11.4% inter-assay %CV) and 0.10 ng/mL for 11-dehydrocorticosterone (5.6% intra-assay, 6.0% inter-assay %CV) and recovery through supported liquid extraction SLE200 plate, was 111% for aldosterone, 118% for corticosterone and 118% for 11-dehydrocorticosterone at LLOQ.

### Data Analysis

FlexImaging™ data were processed and exported as imzML files and further processed with MSiReader™. MSI maps were superimposed with images of the same sections post-staining to define histological zones. Rectangular regions of interest (ROIs; range 2.03-3.54 mm^2^) were drawn in the central area of each kidney section, from pelvic region to outer cortical edges. Within selected ROIs, signal intensities of ions within individual pixels for derivatives of corticosterone, 11-dehydrocorticosterone and aldosterone and their internal standards were exported into MATLAB and processed using an algorithm (Supplementary File 1), which calculated the average of the signal intensity of pixels belonging to the same column of the ROI. These values were plotted as histograms (Fig 2) indicating profiles of steroids of interest. Spatial alignment of histological areas was performed through data imputation to render datasets spatially comparable between different kidney sections. The number of columns for each kidney zone was defined in the widest section. Histological areas were defined in all kidney sections based on histological images and markers. Sections that had <70% of the number of columns per zone were subject to data imputation, with imputed points distributed equally. Imputed values were the average of the adjacent data points with a maximum of 20% of imputed points included.

Untargeted markers of histology were assessed using FlexImaging™, and Lipostar™. Histological features were used to define ROIs and candidate biomarkers that localised to these regions identified.

## Results and Discussion

Steroid hormones ionise poorly and previously derivatisation by Girard T was successful in enhancing signals such that steroids could be mapped on tissue surfaces including brain and testes (13–15). The approach was modified successfully, here (Figs 1Ai, Bi and Ci) as follows, to enhance signal and minimise analyte diffusion. Kidneys were optimally sectioned at −20 °C. Matrix application was performed by automatic sprayer (density of 4 μg/mm^2^) and this approach was found more reproducible than previous manual spraying. Incubation of kidney sections during derivatisation openly in a humidity oven (75%) gave strongest signal intensities and least diffusion (Supplementary File 2). For the purposes of MALDI imaging only one internal standard was used D_8_-corticosterone. In our previous study (13), we had demonstrated that steroids sharing an A ring with similar chemistry to corticosterone reacted with similar efficiency and thus yielded a similar signal intensity due to the ions being generated through the quaternary ammonium function of the derivative. The intensity of signal for derivatised aldosterone (in standard spots) was however lower than that of corticosterone (1.53 × 10^9^ cps vs 11-dehydrocorticosterone (2.65 × 10^9^ cps), and aldosterone (1.03 × 10^9^ cps)). Future inclusion of D_8_-aldosterone may allow for improved quantitation but for this initial proof of principle experiment, the large amounts required to be dissolved in the matrix solution present an economic challenge.

**Figure 1:**
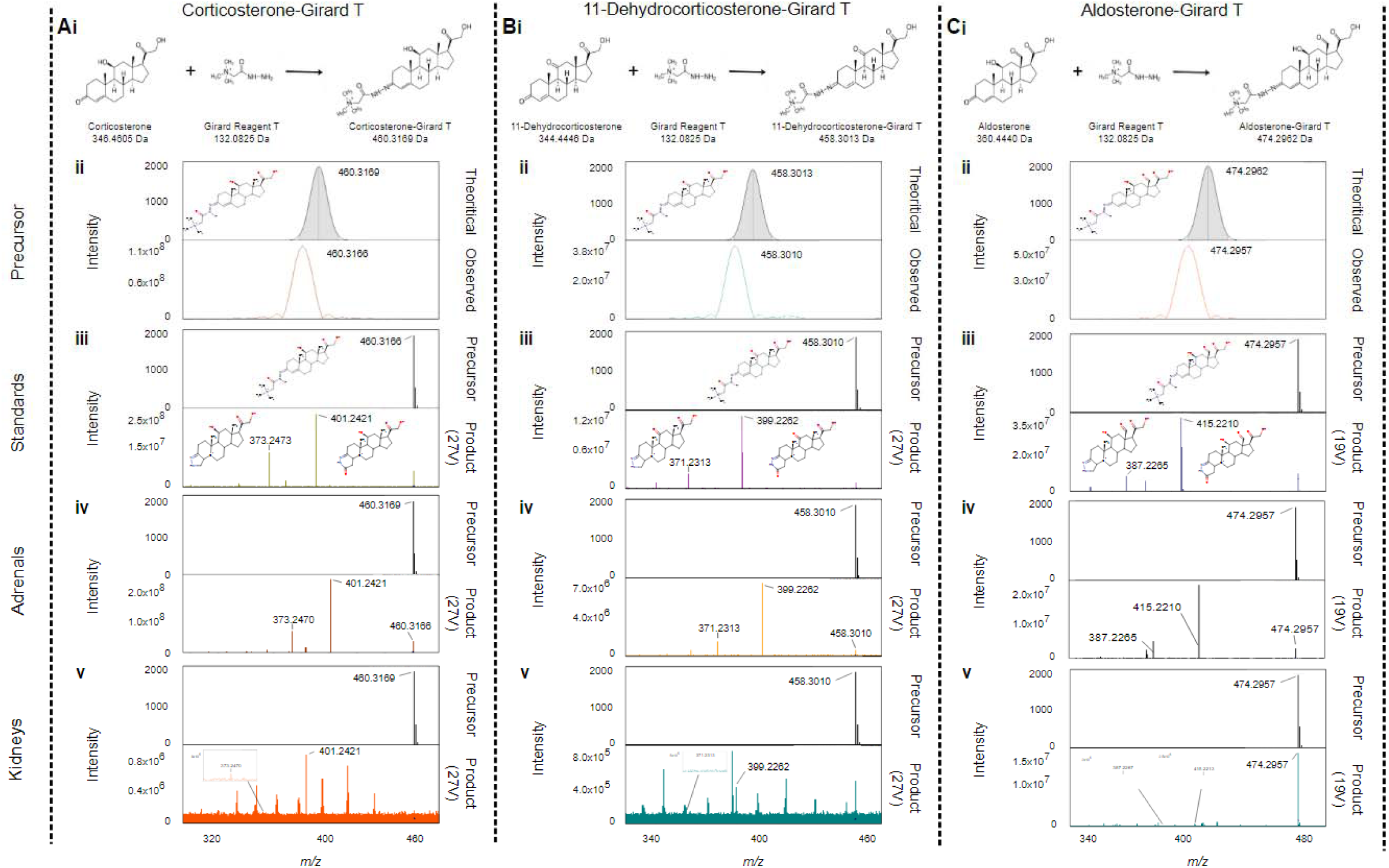
Formation of Girard T derivatives of (Ai) corticosterone, (Bi) 11-dehydrocorticosterone and (Ci) aldosterone. Mass spectra measured by matrix assisted laser desorption ionisation Fourier Transform ion cyclotron resonance mass spectrometry yielding molecular ions with *m/z* 460.3166, 458.3010, and 474.2957 respectively (Aii, Bii, Cii), with mass accuracy of Δppm 0.65, 0.65 and 1.05 from their theoretical masses. Fragmentation spectra were obtained from precursor ions of Girard T derivatives of analytical standards of steroids subject to collision induced dissociation of 27, 27 and 19 eV respectively (Aiii, Biii,Ciii) following LESA sampling. Comparable fragmentation spectra of steroid derivatives were obtained from surfaces of rat adrenal gland (Aiv, Biv, Civ) and mouse kidneys (Av, Bv, Cv).

**Figure 2:**
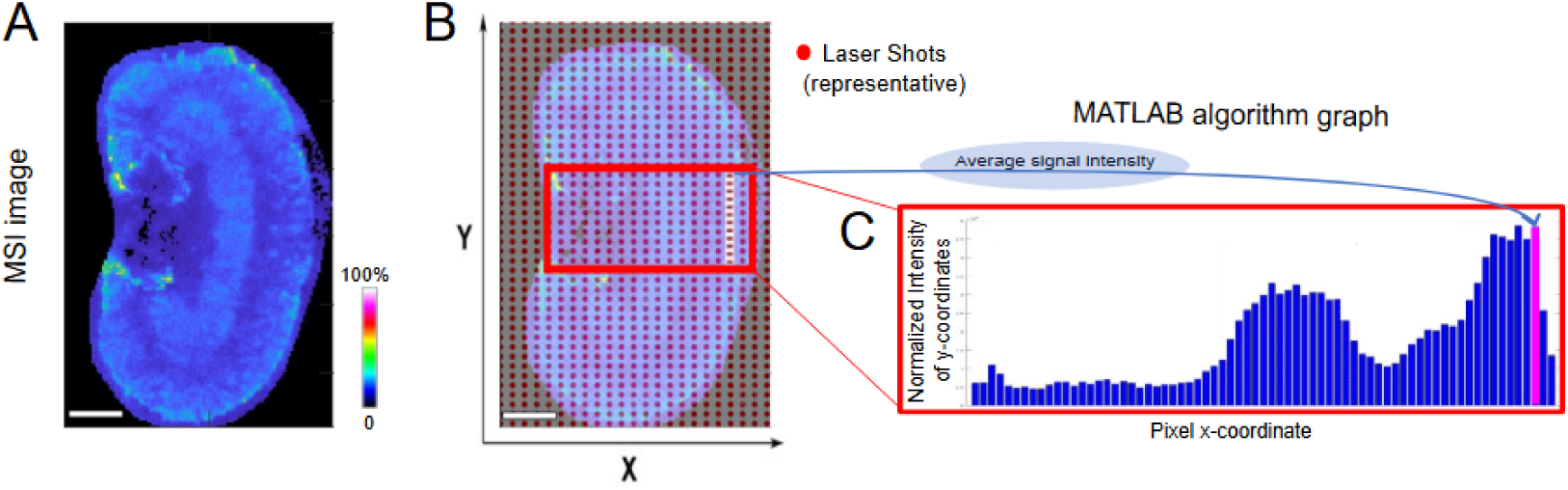
(A) Tissue cryosections were sampled using a (B) rastered laser pattern (red dots show subset of laser shots). Average abundances of data pertaining to each pixel were collated across a rectangular area using a MATLAB algorithm (representative ion *m/z* 476.308). Histograms were reconstructed of relative abundances normalised to intensity of the internal standard (D_8_-Corticosterone) signal within the same pixel. Size alignment bar = 1 mm. MSI= mass spectrometry image.

### Detection and identification of steroids in kidney sections

Mass spectra of GirT derivatives of corticosterone, 11-dehydrocorticosterone and aldosterone in kidney sections yielded base peaks of the molecular ions of *m/z* 460.3166, 458.3010, and 474.2957 sufficiently close to their theoretical *m/z* 460.3169 (Δppm=0.65), 458.3013 (Δppm=0.65) and 474.2962 (Δppm=1.05) respectively (Figs1 Aii, Bii, Cii). Following LESA, the precursor ion of corticosterone-GirT, at *m/z* 460.3166 yielded product ions *m/z* 401.2421 and 373.2473, the precursor ion for 11-dehydrocorticosterone-GirT *m/z* 458.3010 gave product ions *m/z* 399.2262 and 371.2313, and the precursor ion for aldosterone-GirT *m/z* 474.2957 gave product ions *m/z* 415.2210 and 387.2265. (Figs 1 Aiii, Biii, Ciii), typical of fragmentation and rearrangement previously observed for the derivative function of the steroid moiety on the A ring (13). Derivatives of steroids in rat adrenal sections as positive controls also fragmented similarly (Figs 1Aiv, Biv, Civ), within the zona glomerulosa (site of aldosterone synthesis) and zona fasciculata (corticosterone synthesis). In kidney, the ions detected matched those of the standards and fragmentation patterns were in agreement (Figs 1Av, Bv, Cv).

### Distribution of steroids in kidney sections

Histograms of intensities of signals across the rectangular ROIs (Fig 2) were derived to profile the relative quantities of steroids of interest. These were presented as ratios of endogenous steroid to D_8_-corticosterone GirT (internal standard) to account for regional variable ion suppression across tissues and matrix inhomogeneity. To achieve this, sections were stained post-MALDI to identify major renal zones (pelvis, inner/outer cortex and medulla; Fig 3B). This improved alignment over adjacent sections. Spatial alignment of steroid profiles by this approach was complex using histological features alone and further structural detail was sought using integral ion markers.

**Figure 3:**
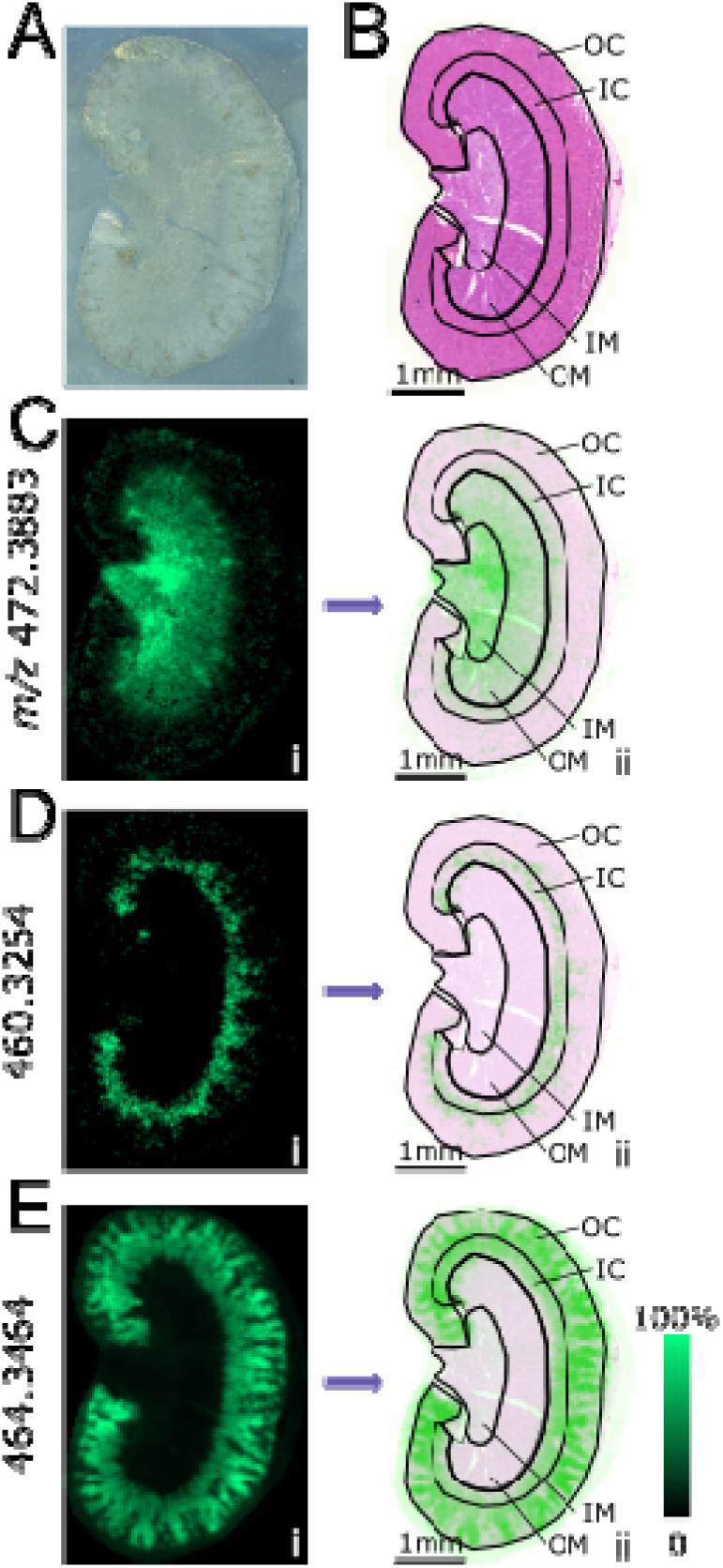
Biomarkers of renal histology. Untargeted datasets collected by mass spectrometry imaging of kidney sections from male C57Bl6/j mice maintained on control diet (0.3% sodium chloride) show structural patterns indicative of histological zones (A) optical scan, (B) adjacent section stained with haematoxylin and eosin, and ions depicting the (Ci) inner (IM) and outer medulla (OM), (Di), inner (IC) and, (Ei) outer cortex (OC)) overlaid with histological zones (ii).

### Mapping renal features using untargeted ion markers

MSI datasets contained many ions beyond the targeted steroids with patterns corresponding to potential nephron segments. These markers allowed integral alignment with histological zones and prevented errors due to changes in shape due to staining. The dataset was analysed first by unsupervised clustering but this had varying success. Thus a supervised approach was adopted whereby regions were approximated manually, inferred from histologically staining (post-imaging) and then ions aligned. This yielded a series of representative ions depicting features of renal architecture (Figs 3C, D, E), allowing more confident identification of medullary/cortical junctions.

Maps thus constructed showed that corticosterone was widely spread, and particularly localised in papilla, medulla and inner cortex (Figs 4B, E, I), while 11-dehydrocorticosterone was more concentrated in the medulla (Fig 4C, F, J). Aldosterone was found in higher amounts in medulla and outer cortex (Figs 4D, G, K). Steroids have not been previously localised in kidney by MSI although binding of radiolabeled corticosterone was localised by autoradiography (12). It might be expected that levels of 11-dehydrocorticosterone would be highest in areas richly expressing 11βHSD2, chiefly in principal cells in the cortical and outer medullary collecting duct. Indeed, areas showing high levels of aldosterone and 11-dehydrocorticosterone, such as inner medulla, could be suggested as areas where MR signaling is primarily regulated by aldosterone. MR regulated processes in this area include stimulation of Na^+^/K^+^-ATPase, localised to the basolateral membrane of most nephron cell types, increasing electrochemical drive for Na^+^ reabsorption in epithelial cells (17) and regulation of ENaC trafficking to prolong the half-life of channel complexes in the apical membrane. These immediate (<4 h) regulatory actions rapidly stimulate epithelial sodium transport. Longer-term, aldosterone activation of MR also increases expression of Na^+^/K^+^-ATPase and ENaC subunits, as well as ROMK potassium channels. The presence of corticosterone was most marked in the outer medulla, where GR is expressed as well as MR. (17) GR activation is now recognised to stimulate sodium and calcium transport in this region, activating thiazide-sensitive NaCl cotransporters (18, 19) and increasing expression of apical calcium channels and binding proteins. (20)

**Figure 4:**
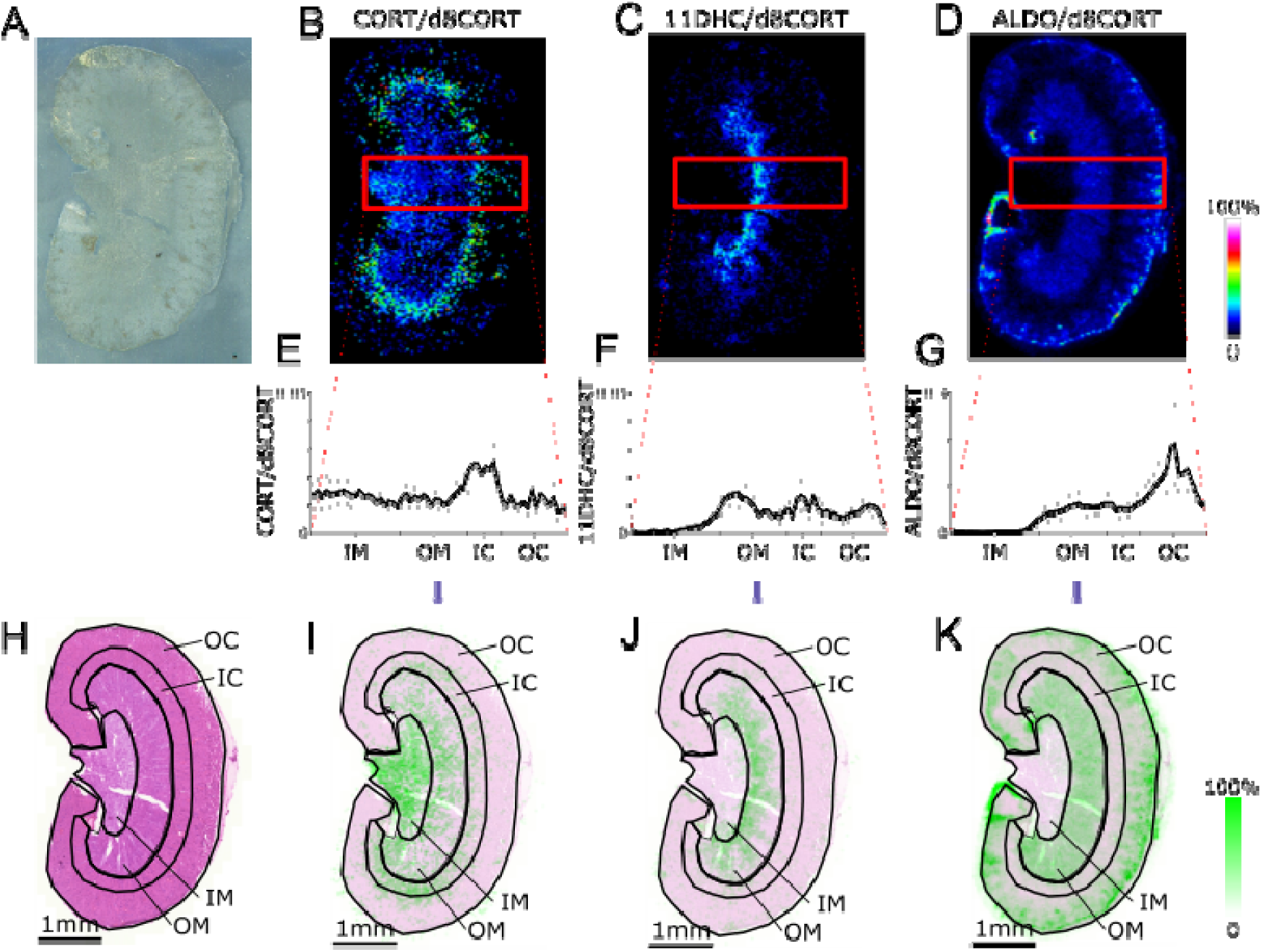
Mass spectrometry images of cryosections of kidneys from male C57Bl6/j mice. (age 12 weeks) maintained on control diet (0.3% sodium chloride) using matrix assisted laser desorption ionisation (75 μm raster X x Y µm) coupled to Fourier transform ion cyclotron resonance mass spectrometry. (A) Optical image of kidney sections prior to analysis. Representative maps of distribution of targeted ions pertaining to Girard T derivatives of (B) corticosterone (CORT; *m/z* 460.3166), (C) 11-dehydrocorticosterone (11DHC; *m/z* 458.3010) and (D) aldosterone (ALDO; *m/z* 474.2957) normalised to D_8_-corticosterone (D_8_-CORT, *m/z* 468.3668). Scale bar = 1 mm and intensity of ion signal shown against a color scale. Data within a rectangular region were normalised against D_8_-corticosterone as an internal standard sprayed across the tissue in conjunction with the derivatisation reagent. Histograms (mean ± standard error of the mean, n=6) are shown for (E) corticosterone, (F) 11-dehydrocorticosterone and (G) aldosterone corrected for internal standard. Steroid ion maps were superimposed with (H) images from adjacent sections stained with hematoxylin and eosin showing that (I) corticosterone localised in the inner medulla (IM), the inner part of the outer medulla (OM) and inner cortex (IC). (J) 11-dehydrocorticosterone was mainly in the inner part of the outer medulla and (K) aldosterone in the outer medulla and outer cortex (OC) (K).

Glucocorticoids are secreted into blood in circadian and ultradian rhythms and thus the amounts of these steroids in the kidney may vary according to these rhythms. The rate at which steroid levels change in kidney will reflect a number of parameters including physicochemical properties of the steroid and tissue and transport in and out of cells. In our previous studies, using tracer infusions to assess tissue penetration, the equilibration rates of tracer steroids varied considerably between liver, adipose and brain (21, 22). The equilibration kinetics of glucocorticoid in healthy liver, with its substantial blood flow and lack of lipid droplets, showed close alignment with blood (22) and kidney may be similar but this should be studied further and the data here recognised as reflecting a single snapshot in time.

Dietary salt stimulates the endocrine regulatory loops of the renin-angiotensin-aldosterone system crucially to homeostatically correct fluid resorption and thus blood pressure. Most knowledge of this axis comes from study of circulating hormones where aldosterone levels are considered key but MSI adds value in unpicking changes in steroids in the tissue setting (Fig 6). Recent studies in humans have reported a direct relationship between urinary sodium excretion and urinary cortisol excretion (23, 24). In mice, high salt increases the forward drive of the hypothalamic-pituitary-adrenal axis to synthesise more glucocorticoid. This is detected as a higher concentration in blood and increased excretion in the urine over a 24h period. (25) In the current study, dietary salt caused the anticipated lowering of aldosterone in blood (Fig 5A), reflecting physiological suppression of the renin angiotensin aldosterone system. However, there was no increase in plasma corticosterone (Fig 5B), which is likely due to collection of samples in the morning, when glucocorticoid levels in mice are at the nadir and no effect of salt diet can be observed (25). In contrast, glucocorticoid amounts were higher in kidney tissue with low salt diets, particularly the outer cortex, where it might influence reabsorption of sodium processes through both GR and MR in distal nephron. The intensities of aldosterone signal within kidney tissue were unaffected by dietary salt, with interesting ramifications for the relative concentration of corticosterone versus aldosterone at critical epithelial sites of sodium reabsorption. Corticosterone concentration ranges from 10 to 100 times higher than aldosterone in plasma, but in some kidney areas aldosterone was present in an order of magnitude higher than the glucocorticoids, likely affecting receptor occupancy. Interestingly the inert metabolite, 11-dehydrocorticosterone, reduced in amount in blood when salt was both lowered and raised in the diet (Fig 5C) which might suggest greater availability of active steroid at critical sites, but this was not reflected in tissue. Physiologically, the interplay between aldosterone and glucocorticoid signaling in sodium-transporting epithelial cells is not fully understood (8, 17) Our findings emphasise the complexity and highlight the need for studies to fully understand how the steroid hormone microenvironment is integrated to regulate renal salt transport. Future advances in instrumentation should allow imaging with cellular resolution to unpick the roles of the availability of ligands for MR and GR in distinct tubular types. It must be noted that only male mice were studied due to the primary aim being to evaluate the technology advance but steroid distribution in kidney should be studied next in female mice.

**Figure 5:**
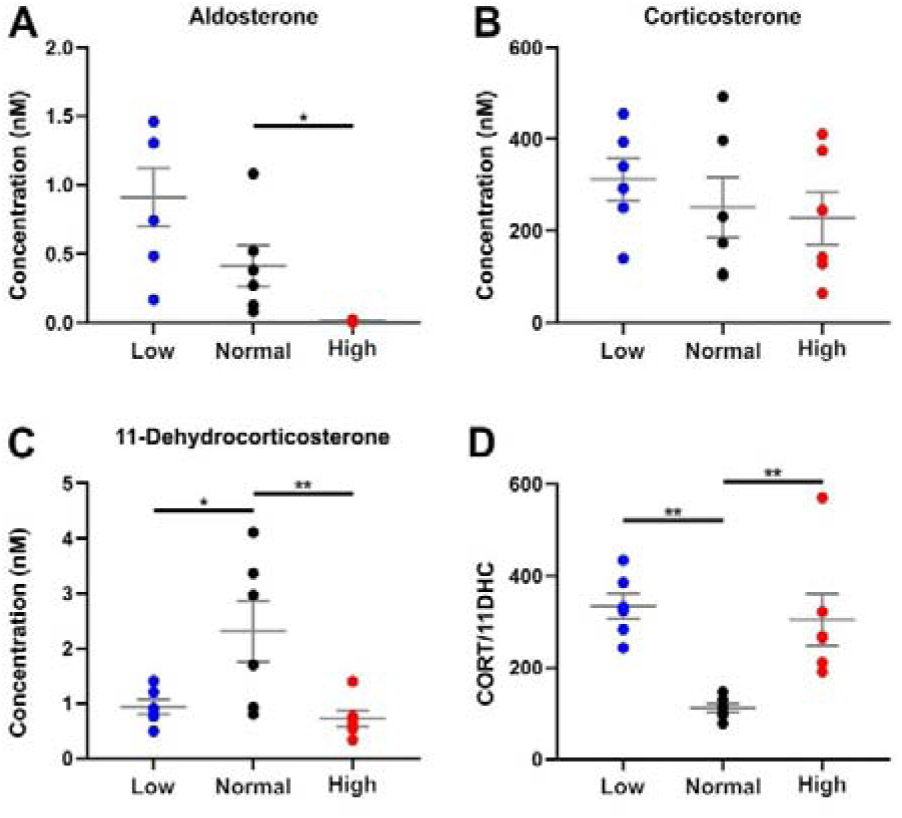
Concentrations of steroids measured in plasma from mice receiving high (3% Na+), control (0.3% Na+) and low (0.03% Na+) salt diets using liquid chromatography tandem mass spectrometry. (A) Concentrations of aldosterone decreased in plasma when salt intake increased. (B) Corticosterone (CORT) concentrations were unaffected when sodium intake was changed, while (C) 11-Dehydrocorticosterone (11DHC) concentrations decreased with both low and high salt intake (D). The ratio of concentrations of active to inactive glucocorticoids was ∼2-fold higher with both low and high salt intake (D). Data are mean = +/−SEM. N=6 compared by ANOVA with Dunnett’s post-hoc tests. * p<0.05, ** p<0.01.

**Figure 6:**
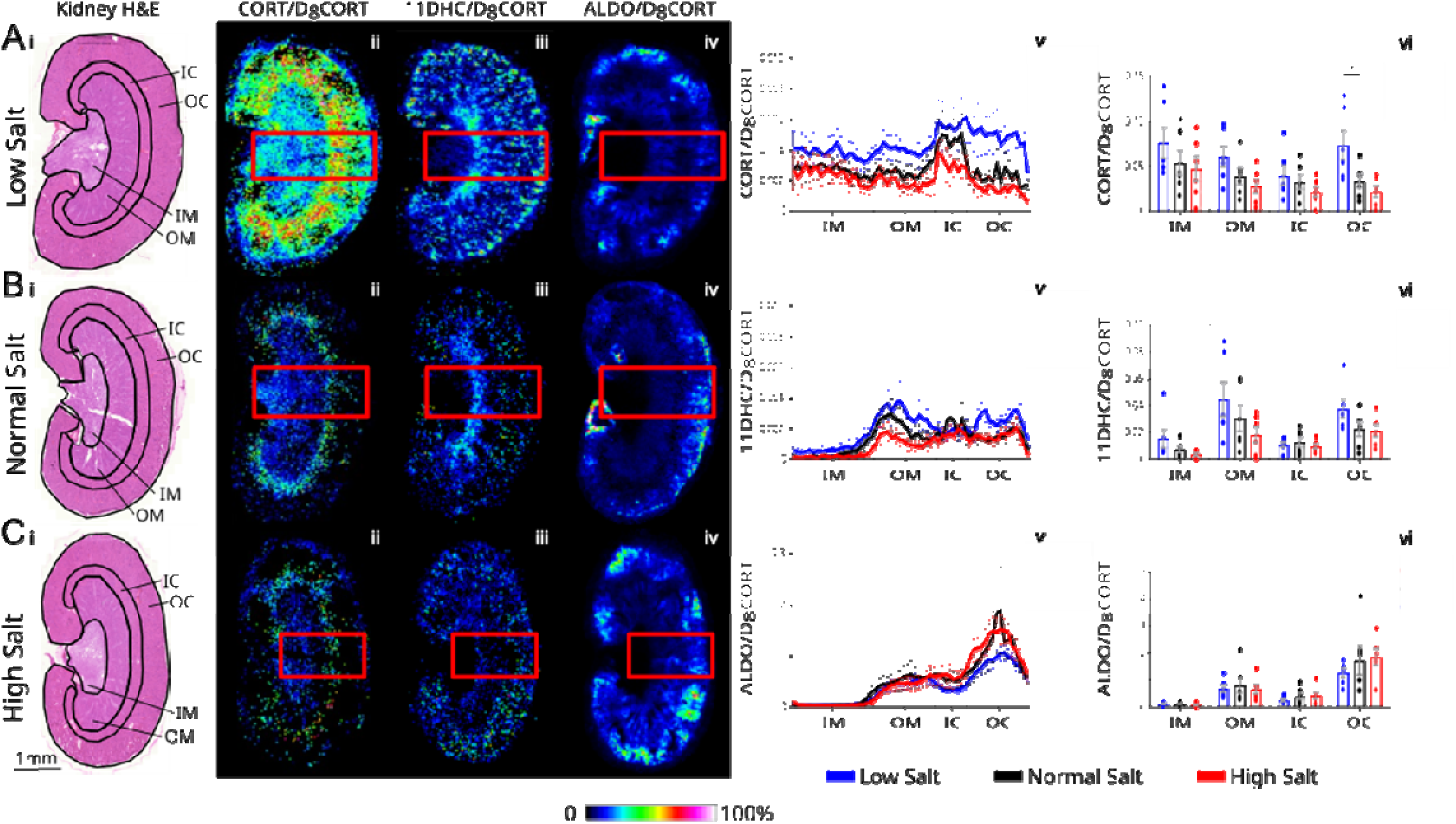
Distribution of steroids in renal cryosections following diets which lower and raise salt intake. Representative histological images of cryosections (adjacent) stained with hematoxylin and eosin (H & E; A, B, Ci) and maps of ions pertaining to Girard T derivatives of corticosterone (CORT; *m/z* 460.3166; A,B,Cii), 11-dehydrocorticosterone (11DHC; *m/z* 458.3010; A,B,Ciii) and aldosterone (ALDO; *m/z* 474.2957; A,B,Civ) normalised to D_8_-corticosterone (D_8_CORT, *m/z* 468.3668), collated from cryosections of kidneys from male C57Bl6/j mice maintained on diets containing low (0.03%), control (0,3%) and high (3%) salt. Scale bar = 1 mm and intensity of ion signal as a color scale. Data were collected using matrix assisted laser desorption ionisation (75 µm raster) coupled to Fourier transform ion cyclotron resonance mass spectrometry. Data are presented as ratios against D_8_-corticosterone as internal standard, sprayed in conjunction with the derivatisation reagent. Ion abundances of steroids (corrected for internal standard) from kidneys of n=6/group are shown in A,B,Cv. Mean values averaged across histological zones are shown in A, B, Cvi. Data are mean±standard error compared by one way ANOVA with Dunnett’s post-hoc test. IM = inner medulla; OM = outer medulla; IC = inner cortex; OC =−outer cortex.

There are several limitations to this study. MS imaging is a powerful technique that offers semi quantitative data and does not allow absolute quantitation. However the use of ratios vs internal standards is a valid approach, as it is considered best practice in other mass spectrometry applications. Quantitation of aldosterone and other steroids of interest could be further improved through use of matched internal standards where available in future work. Furthermore, some steroid metabolites (e.g., conjugates) will not be detected in positive mode, nor will not react with Girard T to produce easily ionised derivatives. Of note renal tissues contain enzymes which catalyse deconjugation and thus this conjugated pool may be of relevance, albeit studied in more detail with DHEA vs glucocorticoids (26, 27). The instrument used in this study allowed spatial resolution down to 50 µm which did not achieve cellular imaging, but importantly instrument advances in the field are opening up new horizons in this arena and becoming more widely available. Care was taken to identify the steroids their *m/z* values with high mass accuracy and resolution using FT-ICR and fragmentation, however the possibility of isobaric interferents (with similar fragmentation patterns) should always be considered when investigating a biochemical pathway with many structurally similar species. Again advances in instrumentation (e.g. ion mobility) may enable workflows that offer greater reassurance regarding specificity moving forward. As a proof of principle study (and to establish statistical power), the experiments were conducted only with male animals culled in the morning. Given the opportunities presented, future studies can now be extended to include wider time frames, both sexes and additional dietary or experimental manipulations. This is of particular importance given the female predisposition to salt-sensitive hypertension (28).

## Conclusions

Application of MSI to understand steroid distribution within renal sections offers new insights into spatial distribution of steroids and thus regions where specific steroids and their cognate receptors functions are likely to dominate. Moreover, MSI shows that plasma values do not report what is happening in tissue which has important ramifications for understanding hormonal control of sodium transport in the kidney. These new insights were not intuitive and worthy of further study including superimposition against renal functional processes e.g. ion channel activity. Glucocorticoids are now believed to hold increasingly important roles in regulating blood pressure particularly related to dietary salt and techniques such as MSI here will help elucidate underlying mechanisms.

## Declarations of Interest

The authors declare that there is no conflict of interest that could be perceived as prejudicing the impartiality of the research reported.

## Funding Sources

We thank Kidney Research UK (ST/009/20161125), the University of Edinburgh Moray Endowment and Deanery of Clinical Science Funds, the Bioscientifica Trust, Society for Endocrinology and the British Heart Foundation (RE/18/5/34216). MR was funded by Wallonia-Brussels International.

## Author Contribution Statement

Conceptualisation (IS, DC, RA, MB, RB); Data curation (IS, SM, NZM, RA); Formal analysis (IS, NZM); Funding acquisition (MB, RA); Investigation (IS, SK, SM, NZM, CLM); Methodology (IS, SK, SM, NZM, DC, CLM); Supervision (RA, MB, RB); Writing – original draft (IS, RA); Writing – review and editing (All authors)

## Acknowledgements

We thank the Edinburgh Clinical Research Facility MS Core (RRID:SCR_021833) and the Little France 2 Biological Research Facility. For the purpose of open access, the author has applied a Creative Commons Attribution (CC BY) license to any Author Accepted Manuscript version arising from this submission.

## Data Availability Statement

Prior to final acceptance, data will be made available on a publicly accessible repository. Matlab Code is available in the Supplementary material.

## Supplementary Information

**File 1:** Coding for Matlab algorithm, software v.R2022a.

**File 2:** Method development in kidney to adapt the method of Cobice et al (13). The main change was incubation was performed in a humidity oven under open conditions and not a sealed container. Success of derivatisation was assessed by applying a spot of D_8_ corticosterone standard (0.5 ng) on the surface of a cryosection of kidney. Derivatisation with Girard T (GirT) and matrix application were performed and intensities and localisation of the ions comprising the spot assessed using matrix assisted laser desorption ionisation Fourier Transform ion cyclotron resonance mass spectrometry. Data were extracted as pixels and distributions represented in histograms, assessed for normal distribution patterns by Kolmogorov-Smirnov (KS) tests. (h=1 if the tests reject the null hypothesis that the dataset follows normal distribution at a 5% significance level, and “0” otherwise; red line=normal distribution trend line). Higher signal intensity and less diffusion was observed when humidity was 75% (C). Scale bar = 1 mm and intensity of ion signal shown against a color scale. Cnts = counts; ROI = region of interest. CHCA = α-cyano-4-hydroxycinnamic acid.

## Notes

### Competing Interest Statement

The authors have declared no competing interest.

