## Supplementary File 1 for "Localisation of corticosteroids in male mouse kidney by mass spectrometry imaging"

**Supplementary File 1: MATLAB algorithm for calculation of signal intensity on kidney sections**

clc

clear all

close all

load FILE_NAME_A_GirT.txt

load FILE_NAME_B_GirT.txt

load FILE_NAME_Aldo_GirT.txt

load FILE_NAME_d8B_GirT.txt

load FILE_NAME_CHCA.txt

load FILE_NAME_mz_4603264.txt

load FILE_NAME_mz_4643464.txt

load FILE_NAME_mz_4723862.txt

[intensity_a, a_coord] = compute_average_intensity( FILE_NAME_A_GirT(:,2), FILE_NAME_A_GirT(:,5) );

[intensity_b, b_coord] = compute_average_intensity( FILE_NAME_B_GirT(:,2), FILE_NAME_B_GirT(:,5) );

[intensity_al, al_coord] = compute_average_intensity( FILE_NAME_Aldo_GirT(:,2), FILE_NAME_Aldo_GirT(:,5) );

[intensity_d8b, d8b_coord] = compute_average_intensity( FILE_NAME_d8B_GirT(:,2), FILE_NAME_d8B_GirT(:,5) );

[intensity_ch, ch_coord] = compute_average_intensity( FILE_NAME_CHCA(:,2), FILE_NAME_CHCA(:,5) );

[intensity_mz460, mz460_coord] = compute_average_intensity( FILE_NAME_mz_4603264(:,2), FILE_NAME_mz_4603264(:,5) );

[intensity_mz464, mz464_coord] = compute_average_intensity( FILE_NAME_mz_4643464(:,2), FILE_NAME_mz_4643464(:,5) );

[intensity_mz472, mz472_coord] = compute_average_intensity( FILE_NAME_mz_4723862(:,2), FILE_NAME_mz_4723862(:,5) );

figure(1)

A = bar(a_coord, intensity_a);

A.FaceColor = 'b';

hold on

xline (46.5000,'--r',{'Renal pelvis'});

xline (48.1125,'--r',{'Medullary edge'});

xline (49.2375,'--r',{'Corticomedullary',' juction'});

xline (49.7625,'--r',{'Inner cortical edge'});

xline (50.8875,'--r',{'Outer cortical edge'});

ylabel('Average intesity','fontsize',16)

title('A-GirT average intensity along X-axis','fontsize',20)

set(gca,'XTick',[(46.5000+48.1125)/2 (48.1125+49.2375)/2 (49.2375+49.7625)/2 (49.7625+50.8875)/2],'XTickLabel', {'Papilla', 'Medulla', 'Inner Cortex', 'Outer Cortex' }, 'FontSize', 20)

print (gcf, 'A-GirT intensity ver2.png','-dpng','-r300');

figure(2)

A = bar(b_coord, intensity_b);

A.FaceColor = 'b';

hold on

xline (46.5000,'--r',{'Renal pelvis'});

xline (48.1125,'--r',{'Medullary edge'});

xline (49.2375,'--r',{'Corticomedullary',' juction'});

xline (49.7625,'--r',{'Inner cortical edge'});

xline (50.8875,'--r',{'Outer cortical edge'});

ylabel('Average intesity','fontsize',16)

title('B-GirT average intensity along X-axis','fontsize',20)

set(gca,'XTick',[(46.5000+48.1125)/2 (48.1125+49.2375)/2 (49.2375+49.7625)/2 (49.7625+50.8875)/2],'XTickLabel', {'Papilla', 'Medulla', 'Inner Cortex', 'Outer Cortex' }, 'FontSize', 20)

print (gcf, 'B-GirT intensity ver2.png','-dpng','-r300');

figure(3)

A = bar(al_coord, intensity_al);

A.FaceColor = 'b';

hold on

xline (46.5000,'--r',{'Renal pelvis'});

xline (48.1125,'--r',{'Medullary edge'});

xline (49.2375,'--r',{'Corticomedullary',' juction'});

xline (49.7625,'--r',{'Inner cortical edge'});

xline (50.8875,'--r',{'Outer cortical edge'});

ylabel('Average intesity','fontsize',16)

title('Aldo-GirT average intensity along X-axis','fontsize',20)

set(gca,'XTick',[(46.5000+48.1125)/2 (48.1125+49.2375)/2 (49.2375+49.7625)/2 (49.7625+50.8875)/2],'XTickLabel', {'Papilla', 'Medulla', 'Inner Cortex', 'Outer Cortex' }, 'FontSize', 20)

print (gcf, 'Aldo-GirT intensity ver2.png','-dpng','-r300');

figure(4)

A = bar(d8b_coord, intensity_d8b);

A.FaceColor = 'b';

hold on

%xline (47.9625,'--r',{'Cortex'});

xline (46.5000,'--r',{'Renal pelvis'});

xline (48.1125,'--r',{'Medullary edge'});

xline (49.2375,'--r',{'Corticomedullary',' juction'});

xline (49.7625,'--r',{'Inner cortical edge'});

xline (50.8875,'--r',{'Outer cortical edge'});

ylabel('Average intesity','fontsize',16)

title('d8B-GirT average intensity along X-axis','fontsize',20)

set(gca,'XTick',[(46.5000+48.1125)/2 (48.1125+49.2375)/2 (49.2375+49.7625)/2 (49.7625+50.8875)/2],'XTickLabel', {'Papilla', 'Medulla', 'Inner Cortex', 'Outer Cortex' }, 'FontSize', 20)

print (gcf, 'd8B-GirT intensity ver2.png','-dpng','-r300');

figure(5)

A = bar(ch_coord, intensity_ch);

A.FaceColor = 'b';

hold on

%xline (47.9625,'--r',{'Cortex'});

xline (46.5000,'--r',{'Renal pelvis'});

xline (48.1125,'--r',{'Medullary edge'});

xline (49.2375,'--r',{'Corticomedullary',' juction'});

xline (49.7625,'--r',{'Inner cortical edge'});

xline (50.8875,'--r',{'Outer cortical edge'});

ylabel('Average intesity','fontsize',16)

title('CHCA average intensity along X-axis','fontsize',20)

set(gca,'XTick',[(46.5000+48.1125)/2 (48.1125+49.2375)/2 (49.2375+49.7625)/2 (49.7625+50.8875)/2],'XTickLabel', {'Papilla', 'Medulla', 'Inner Cortex', 'Outer Cortex' }, 'FontSize', 20)

print (gcf, 'CHCA intensity ver2.png','-dpng','-r300');

function [mean_intensity, unique_coord] = compute_average_intensity(coord, intensity)

unique_coord = unique(coord);

mean_intensity = zeros(length(unique_coord), 1);

for i = 1:length(unique_coord)

I = find(unique_coord(i) == coord);

mean_intensity(i)=mean( intensity(I) )

end

end
