## Supplementary File 2 for "Localisation of corticosteroids in male mouse kidney by mass spectrometry imaging"

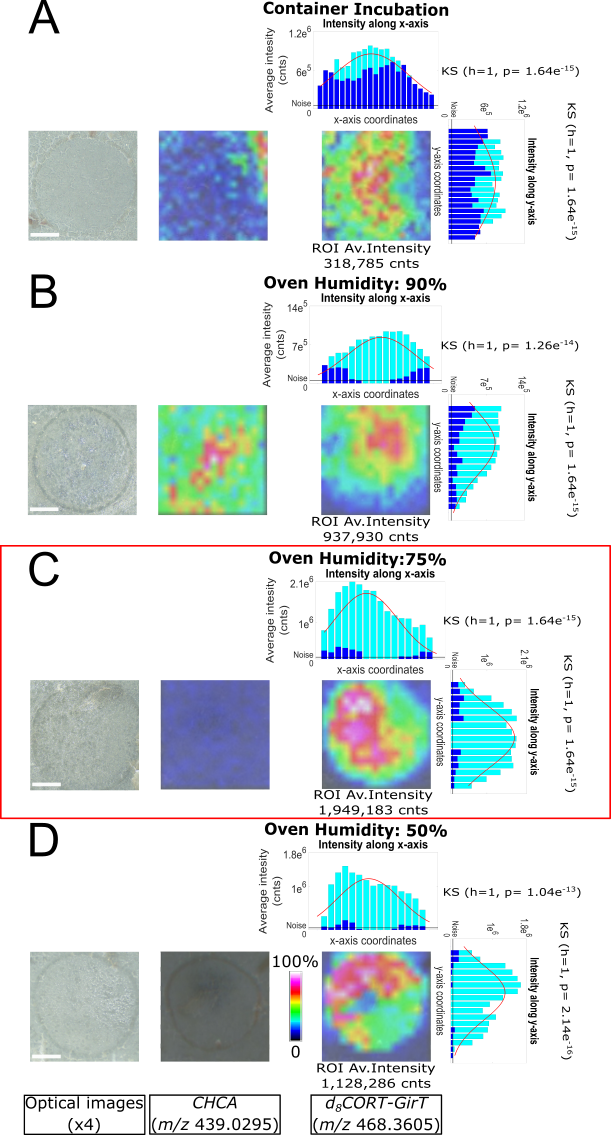

**Supplementary File 2:** Method development in kidney to adapt the method of Cobice et al [13]. The main change was incubation was performed in a humidity oven under open conditions and not a sealed container. Success of derivatization was assessed by applying a spot of D_8_ corticosterone standard (0.5 ng) on the surface of a cryosection of kidney. Derivatization with Girard T (GirT) and matrix application were performed and intensities and localization of the ions comprising the spot assessed using matrix assisted laser desorption ionization Fourier Transform ion cyclotron resonance mass spectrometry. Data were extracted as pixels and distributions represented in histograms, assessed for normal distribution patterns by Kolmogorov-Smirnov (KS) tests. (h=1 if the tests reject the null hypothesis that the dataset follows normal distribution at a 5% significance level, and “0” otherwise; red line=normal distribution trend line). Higher signal intensity and less diffusion was observed when humidity was 75% (C). Scale bar = 1 mm and intensity of ion signal shown against a color scale. Cnts = counts; ROI = region of interest. CHCA = α-cyano-4-hydroxycinnamic acid.

[13] D.F. Cobice, C.L. MacKay, R. Goodwin, A. McBride, P. Langridge-Smith, S.P. Webster, B.R. Walker, R. Andrew, MS imaging for dissecting steroid intracrinology within target tissues., Analytical Chemistry 85 (2013) 11576-11584.
